# Autonomous bioluminescence imaging of single mammalian cells with the bacterial bioluminescence system

**DOI:** 10.1101/798108

**Authors:** Carola Gregor, Jasmin K. Pape, Klaus C. Gwosch, Tanja Gilat, Steffen J. Sahl, Stefan W. Hell

## Abstract

Bioluminescence based imaging of living cells has become an important tool in biological and medical research. However, many bioluminescence imaging applications are limited by the requirement of an externally provided luciferin substrate and the low bioluminescence signal which restricts the sensitivity and spatiotemporal resolution. The bacterial bioluminescence system is fully genetically encodable and hence produces autonomous bioluminescence without an external luciferin, but its brightness in cell types other than bacteria has so far not been sufficient for imaging single cells. We coexpressed codon-optimized forms of the bacterial *luxCDABE* and *frp* genes from multiple plasmids in different mammalian cell lines. Our approach produces high luminescence levels that are comparable to firefly luciferase, thus enabling autonomous bioluminescence microscopy of mammalian cells.

**Significance statement:** Bioluminescence is generated by luciferases that oxidize a specific luciferin. The enzymes involved in the synthesis of the luciferin from widespread cellular metabolites have so far been identified for only two bioluminescence systems, those of bacteria and fungi. In these cases, the complete reaction cascade is genetically encodable, meaning that heterologous expression of the corresponding genes can potentially produce autonomous bioluminescence in cell types other than the bacterial or fungal host cells. However, the light levels achieved in mammalian cells so far are not sufficient for single-cell applications. Here we present, for the first time, autonomous bioluminescence images of single mammalian cells by coexpression of the genes encoding the six enzymes from the bacterial bioluminescence system.

---

Bioluminescence is an enzyme-catalyzed process by which living cells can produce light. It requires a luciferin substrate that is oxidized by a luciferase, a reaction during which the bioluminescence light is emitted. In addition to the luciferase, several other enzymes participate in the process of bioluminescence by synthesizing the luciferin from cellular metabolites and by converting it back into its reduced form. For most bioluminescence systems, these proteins have not fully been identified so far, with the exception of the bacterial (1–5) and the recently solved fungal system (6). Knowledge of all enzymes participating in the reaction cascade means that they can in principle be heterologously expressed in any cell type in order to produce autonomous bioluminescence for imaging applications. This circumvents the problems associated with external luciferin supply, such as elevated background signal due to non-enzymatic luciferin oxidation in solution (7) or decay of the signal over time upon luciferin consumption (8, 9).

Of particular interest for numerous applications in medical research (10) is the bioluminescence imaging of mammalian cells. However, neither the bacterial nor the fungal bioluminescence system are widely used for this purpose. In the case of fungal bioluminescence, autonomous bioluminescence imaging of mammalian cells has not been demonstrated so far. One obstacle for this is that fungal luciferase loses its activity at temperatures above 30 °C (6). The bacterial bioluminescence system has been implemented in mammalian cells (11, 12), but the resulting bioluminescence is much dimmer than that produced by other luciferases which require an external luciferin (13). The low brightness of bacterial bioluminescence in mammalian cells impedes particularly single-cell imaging applications due to insufficient contrast.

Many scientists in the field have contributed to elucidating the biochemistry of bacterial bioluminescence, which provides the basis for the present work (for reviews, see e.g. (14–18). The light emitter in bacterial bioluminescence is assumed to be FMN-C4a-hydroxide (19, 20) which is formed in its excited state during the bioluminescence reaction and emits a photon upon return to its ground state. During this reaction, FMNH_2_ is oxidized to FMN and a long-chain aliphatic aldehyde (RCHO) is oxidized to the corresponding carboxylic acid (RCOOH):

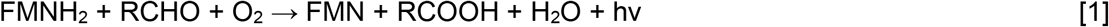

Both products are recycled under consumption of cellular energy. This is performed by a flavin reductase which reduces the oxidized FMN to FMNH_2_ as well as by the fatty acid reductase complex that converts the acid back into the aldehyde:

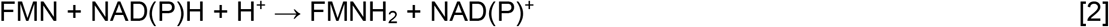

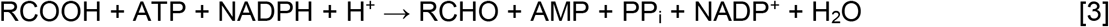

The involved enzymes are coded by the *lux* operon. The genes *luxA* and *luxB* encode the luciferase subunits, whereas *luxC*, *D* and *E* encode the subunits of the fatty acid reductase complex. FMNH_2_ is either produced by the FMN reductase coded by *luxG* or by other cellular enzymes.

Although none of the required substrates is exclusively found in bacteria and hence bacterial bioluminescence should in principle be functional in other cell types as well, expression of the *lux* genes in eukaryotic cells has only been moderately successful so far. It has been demonstrated that codon optimization of the *lux* genes improves their expression in mammalian cells (11, 12, 21), yet the achieved brightness has been comparatively low (12, 13), hampering the prospects for single-cell visualization by microscopy. In addition to variations in codon usage between different cell types, the requirement for simultaneous multi-gene expression of all *lux* proteins at high levels poses further challenges for the use in mammalian cells. Here, we demonstrate that codon-optimized (co) *lux* genes expressed from individual plasmids in an optimized ratio produce bright autonomous bioluminescence in different mammalian cell lines. The resulting signal is comparable to that of firefly luciferase and enables imaging of single cells over extended time periods. Moreover, it allows the observation of toxic compounds effects and starvation on cell viability and metabolic activity.

## Results

### Expression of Lux proteins in mammalian cells

Since the wild-type *lux* genes have been shown to be poorly expressed in mammalian cells in their native form (11), we generated codon-optimized versions of the *luxCDABE* genes from *Photorhabdus luminescens* and the NADPH-flavin oxidoreductase gene *frp* from *Vibrio campbellii* using the codon adaptation tool JCat (http://www.jcat.de, (22)). The resulting genes (Fig. S1) were synthesized, cloned separately into the expression vector pcDNA3.1(+) and cotransfected in equal amounts into HEK 293 cells. While expression of the wild-type genes did not result in detectable bioluminescence levels, the optimized genes produced high signal (Fig. 1). Assuming that a signal threefold higher than the standard deviation of the background (~8 counts per pixel for 10-min exposure time) would have been readily detectable, comparison of the camera counts between wild-type (no signal) and co *lux* (~3000 counts per pixel for 1-min exposure time, Fig. 1) indicated that the gene optimization improved the light output by at least three orders of magnitude. This demonstrates that the Lux proteins are functional in mammalian cells, but that their expression is highly dependent on the codon usage or possibly other factors determined by the nucleotide sequence. Variation of the ratios of the transfected plasmids while keeping the total amount of DNA constant indicated that the brightness was increased by 66% if a threefold molar amount of the fatty acid reductase components *luxC*, *D* and *E* was used (Fig. 2). Therefore, synthesis of the aldehyde seems to be the most rate-limiting step of the overall reaction cascade rather than the luciferase-catalyzed bioluminescence reaction itself.

**Fig. 1.**
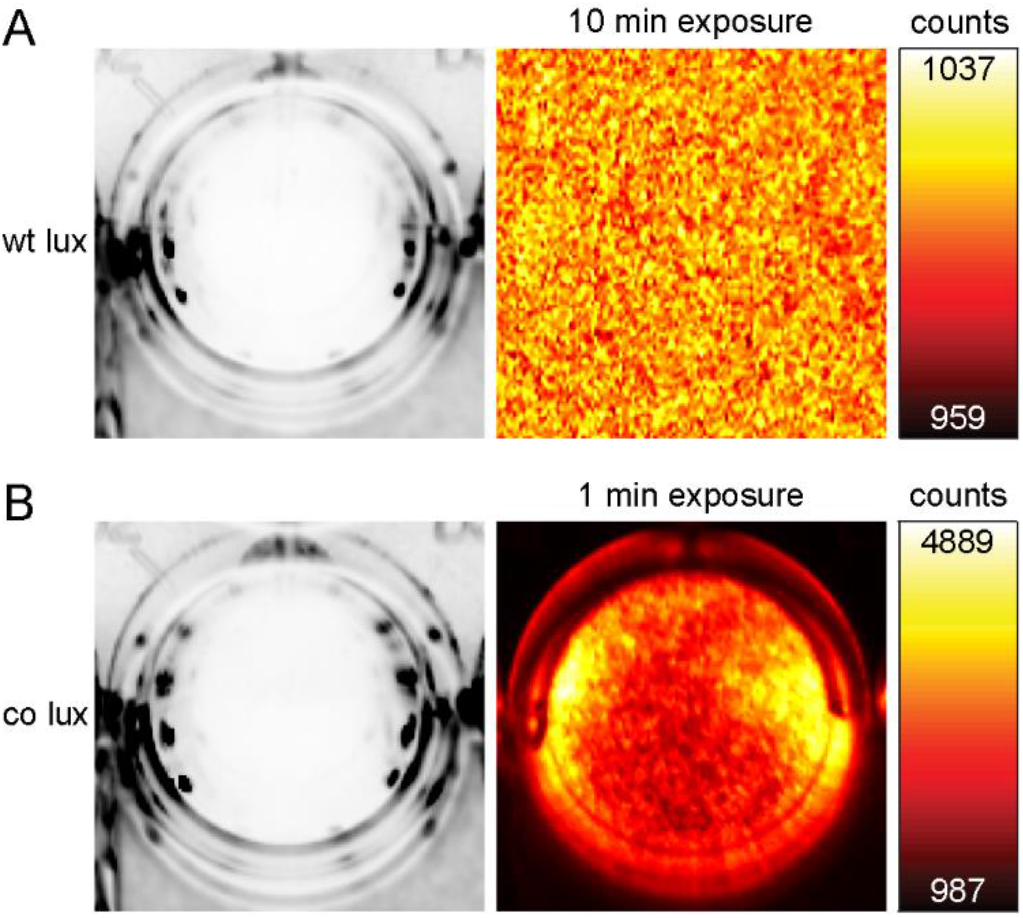
Bioluminescence of wild-type *lux* and codon-optimized (co) *lux*. HEK 293 cells were transfected with wild-type *lux* and *frp* genes without codon optimization (*A*) or the co *lux* genes (*B*) in a ratio of *luxA*:*luxB*:*luxC*:*luxD*:*luxE*:*frp*=1:1:3:3:3:1. Cells were imaged in a 24-well plate with an Amersham Imager 600. Left, white-light images of the respective well with a monolayer of cells, right, bioluminescence signal after the indicated exposure time. The color maps were scaled to the minimum and maximum camera counts per pixel of the bioluminescence images.

Since simultaneous cotransfection of six different plasmids reduces the transfectable amount for each gene, we tested whether the brightness can be increased by reducing the number of plasmids and expressing more than one gene from the same vector. For this purpose, we linked several genes by viral 2A sequences, as this approach has been reported to increase the transfection efficiency and brightness (12). The 2A sequence impairs the formation of a peptide bond during translation and thus results in the synthesis of two separate proteins with the major part of the peptide remaining attached to the C terminus of the first protein (P2A: ATNFSLLKQAGDVEENPG↓P, T2A: EGRGSLLTCGDVEENPG↓P) (23). Although the transfected amount of the combined genes was accordingly increased, none of the tested combinations showed an increase in bioluminescence (Fig. 2). This might be caused by a lower transfection efficiency, decreased translation rates or impaired function of the Lux proteins due to the remaining 2A fragments. Consequently, maximum brightness was achieved when all six proteins were expressed from separate plasmids (Fig. 2).

**Fig. 2.**
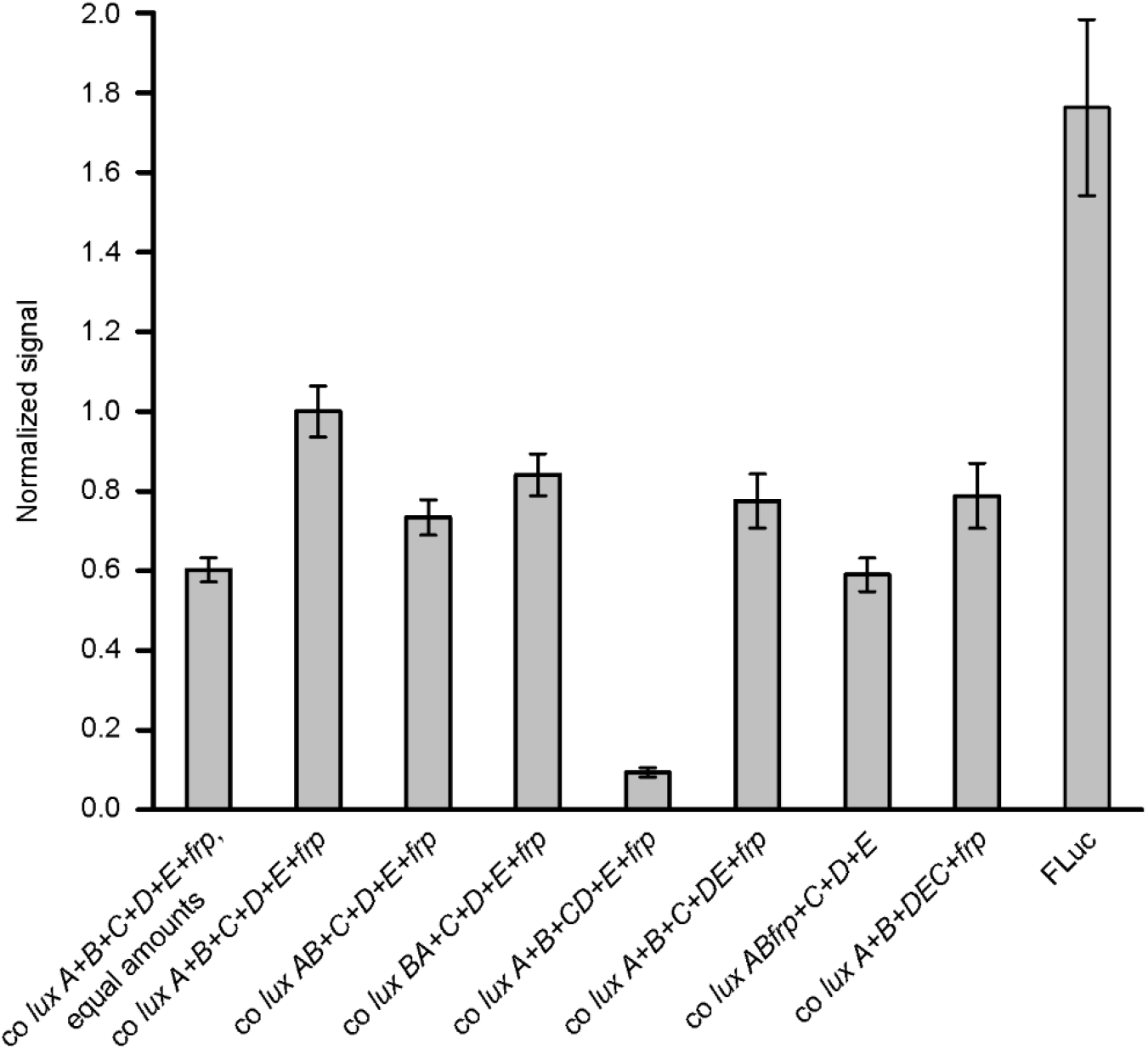
Bioluminescence of HEK 293 cells expressing different combinations of co *lux* genes or firefly luciferase (FLuc) from pcDNA3.1(+). Cells were grown in 24-well plates, transfected with the same total amount of DNA and imaged with an Amersham Imager 600. Unless otherwise stated, plasmids encoding *luxA*, *B*, *C*, *D*, *E* and *frp* were transfected in a ratio of 1:1:3:3:3:1. Plasmids encoding two or three genes were transfected in two- or threefold amount, respectively. The following abbreviations are used: *luxAB*: *luxA*-P2A-*luxB*, *luxBA*: *luxB*-P2A-*luxA*, *luxCD*: *luxC*-P2A-*luxD*, *luxDE*: *luxD*-P2A-*luxE*, *luxABfrp*: *luxA*-P2A-*luxB*-T2A-*frp*, *luxDEC*: *luxD*-P2A-*luxE*-T2A-*luxC*. (+) signs indicate expression from separate plasmids. For imaging of FLuc, 150 μg/mL D-luciferin was added to the medium. Error bars represent SD of the signal of five separate wells.

We compared the brightness of co Lux to the widely used firefly luciferase (FLuc). Although FLuc was transfected in a 12-fold molar amount compared to the *luxAB* luciferase subunits, its brightness was only 76% higher (Fig. 2). Hence, bacterial luciferase is well functional in mammalian cells when efficiently expressed and produces signal comparable to firefly luciferase if cellular production of the substrates is high enough.

Since bioluminescence systems based on the exogenous luciferin coelenterazine often suffer from background signal due to auto-oxidation of coelenterazine in solution, we compared its background to the co Lux system. For this purpose, we measured the signal from non-transfected cells with 5 μM coelenterazine and from cells transfected with the FMNH_2_-producing co *frp* (Fig. S2). Whereas *frp* expression resulted in no detectable bioluminescence emission, non-enzymatic coelenterazine oxidation produced clear signal. Therefore, the low background of the co Lux system allows its bioluminescence detection with a higher specificity than coelenterazine-based systems.

As codon-optimized versions of the *lux* genes have been previously expressed in mammalian cells (11, 12), we compared the brightness of our co Lux system to pCMV_Lux_ (12). pCMV_Lux_ is a humanized *lux* expression plasmid that contains all six *luxCDABE* and *frp* genes separated by 2A sequences and produces the highest levels of autonomous bioluminescence in mammalian cells reported so far. Expression of pCMV_Lux_ in HEK 293 cells was not detectable with our bioluminescence imager at 37 °C (Fig. S3). Weak signal (on average ~20 counts above background in 10 min) was only detectable after the cells had cooled down to room temperature, but the overall brightness was still ~3 orders of magnitude lower than with our co Lux system (Fig. S3, Fig. 1). In order to find out the main reasons for this large difference, we first compared the codon adaptation indices (CAIs) of our co *lux* genes and the humanized genes from pCMV_Lux_ (referred to as h*lux*) and the non-optimized wild-type genes (Table S1). While the low CAIs of the non-optimized genes (average 0.13) may account for poor expression and consequently non-detectable light levels, the difference of the CAIs of the h*lux* and co *lux* genes are smaller (average 0.74 and 0.95, respectively). Therefore, differences in codon usage may not be the only factor that contributes to the decreased brightness from the h*lux* genes. To test the influence of the 2A sequences contained in pCMV_Lux_ on the bioluminescence levels, we cloned the individual h*lux* genes without the 2A sequences into pcDNA3.1(+), analogous to our co *lux* genes. Cotransfection of the h*lux* pcDNA3.1(+) plasmids in HEK 293 cells produced detectable bioluminescence, but only 1–2% of that from the co *lux* pcDNA3.1(+) plasmids (Fig. S4). This suggests that the 2A peptides are one important factor that reduces expression or activity of the Lux proteins, in accordance with the results from our co *lux* 2A constructs (Fig. 2). In addition, we combined single h*lux* pcDNA3.1(+) plasmids with the remaining co *lux* pcDNA3.1(+) plasmids to test the influence of the individual h*lux* genes on bioluminescence levels (Fig. S4). All h*lux* combinations exhibited decreased brightness compared to co *lux*, with the brightness lowest for h*frp*. While the observed differences may partly be attributed to the lower CAIs of the h*lux* genes, other reasons may be responsible for the strongly decreased light emission with the h*frp* gene. The most likely explanation is a lower functionality of the protein in mammalian cells due to multiple differences in the protein sequence since a different host strain was chosen for h*frp* (*V. harveyi* instead of *V. campbellii* for co *frp*). Overall, our co Lux system provides an improvement in brightness of at least three orders of magnitude compared to previous approaches of *lux* gene expression in mammalian cells.

### Test of toxic effects caused by the expression of co *lux*

Since the bacterial bioluminescence reaction requires a long-chain aldehyde substrate that is potentially cytotoxic, we tested if the expression of co *lux* has adverse cellular effects. First, we performed an MTT assay based on the biochemical reduction of the tetrazolium dye 3-(4,5-dimethylthiazol-2-yl)-2,5-diphenyltetrazolium bromide (MTT) by NAD(P)H. Decreased cell growth and viability result in lower formation of the reduced formazan, which can be detected as a lower absorption. HeLa cells transfected with co *lux* or the empty pcDNA3.1(+) vector using Lipofectamine 2000 did not show significant differences (p>0.5) in MTT reduction (Fig. 3*A*). However, lysates of untransfected cells exhibited a 53% higher absorbance, indicating that the only toxicity observed is caused by the transfection reagent rather than expression of the co *lux* genes.

**Fig. 3.**
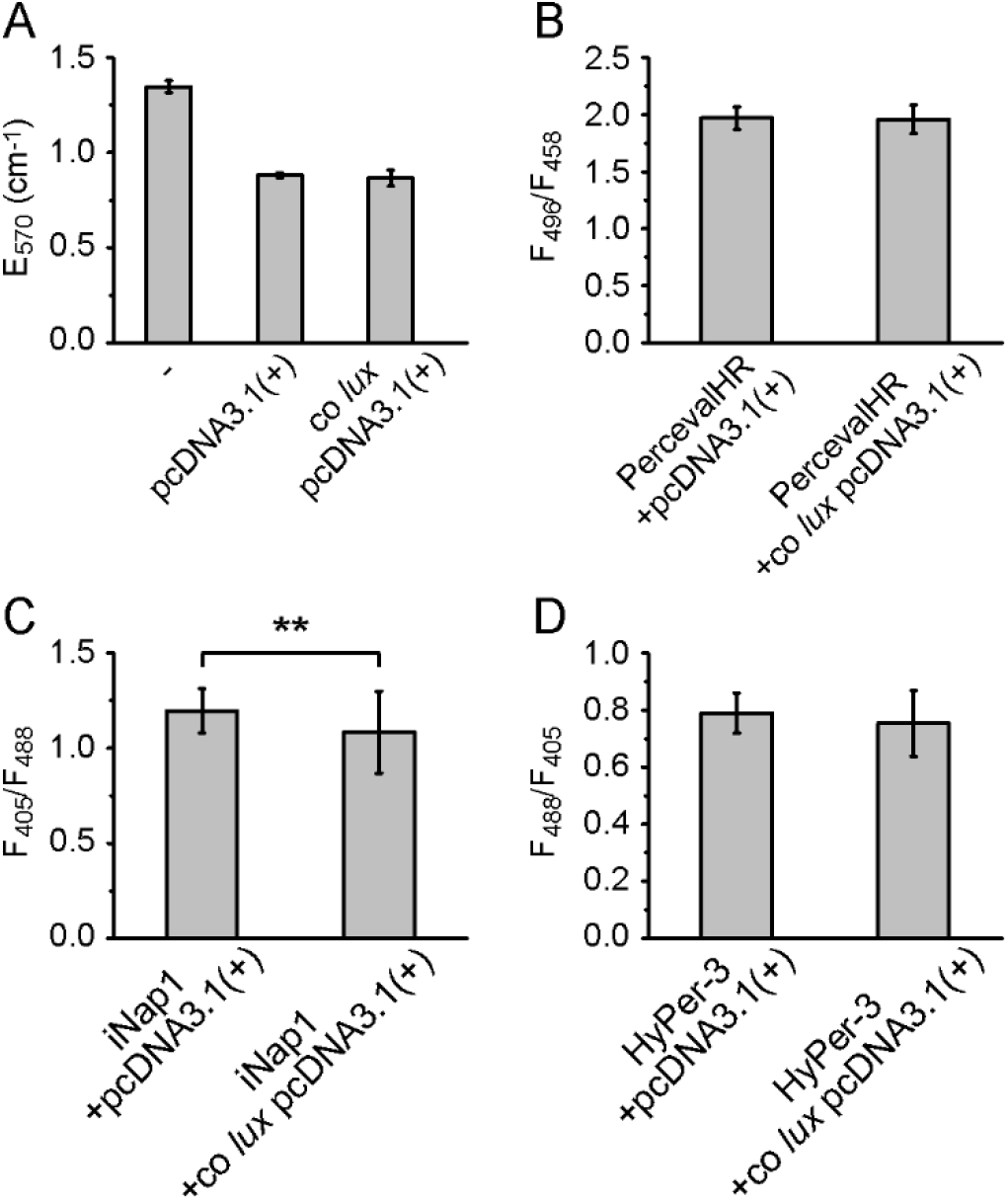
Codon-optimized (Co) *lux* toxicity control measurements. HeLa cells were cotransfected with co *lux* genes in pcDNA3.1(+) or the empty vector, and a fluorescent sensor where indicated. (*A*) Absorbance of cell lysates from MTT assay with untransfected cells (−) for comparison. Error bars represent SD of three separate coverslips. (*B-D*) Comparison of ATP (*B*), NADPH (*C*) and H_2_O_2_ levels (*D*) by ratiometric fluorescence imaging of the sensors PercevalHR (*B*), iNap1 (*C*) and HyPer-3 (*D*) using the indicated excitation wavelengths. Error bars represent SD of 50 cells. ** represents a p-value of <0.01 as calculated by a 2-tailed Student’s *t*-test.

Since toxic effects may involve more subtle cellular changes due to depletion of ATP or NADPH or increased production of hydrogen peroxide due to reaction of the luciferase-bound FMNH_2_ with molecular oxygen (17, 24), we compared their cellular levels with and without co *lux* expression. For this purpose, we cotransfected HeLa cells with a ratiometric fluorescent sensor for ATP, NADPH or H_2_O_2_ and imaged them by fluorescence microscopy (Fig. 4). For imaging of ATP, we used the sensor PercevalHR which measures the cellular ATP-to-ADP ratio. ATP and ADP binding increase its fluorescence excited at 500 nm and 420 nm, respectively; there is a fixed isosbestic point at ~455 nm (25). NADPH concentration was imaged with the sensor iNap1, which increases its fluorescence excited at 420 nm and decreases its fluorescence excited at 485 nm upon NADPH binding (26). H_2_O_2_ levels were compared using the sensor HyPer-3 which increases its fluorescence ratio F500/F420 upon excitation at 500 and 420 nm (27). No significant change of the fluorescence ratios between cells transfected with co *lux* or empty pcDNA3.1(+) vector was observed for ATP and H_2_O_2_ (p>0.05, Fig. 3*B* and *D*). In case of NADPH, we found a small (9%), but significant (p<0.01) change in the iNap fluorescence ratio (Fig. 3*C*). The reason for this might be increased NADPH consumption for the synthesis of the aldehyde or FMNH_2_.

**Fig. 4.**
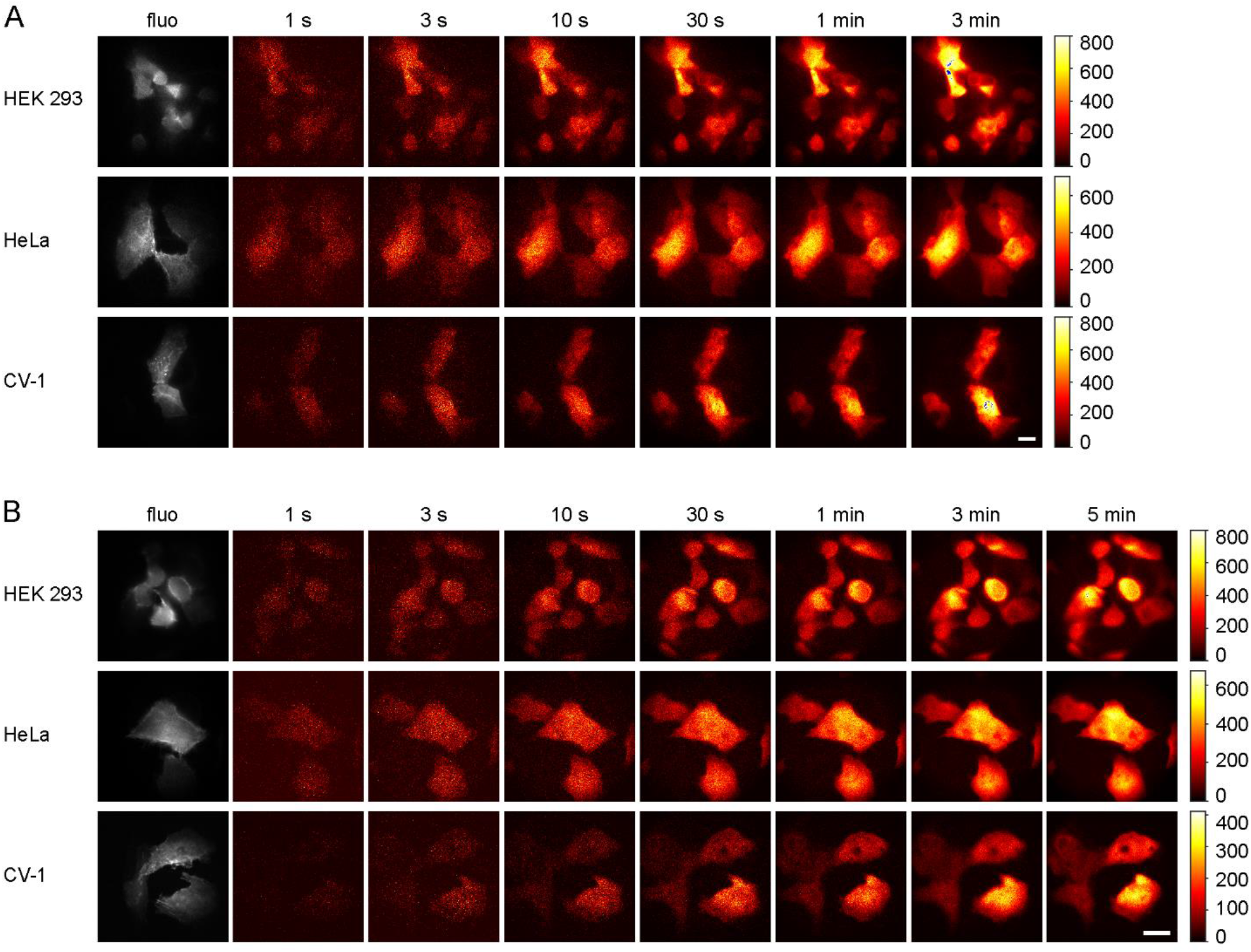
Bioluminescence images of different cell lines expressing the co *lux* genes. Images were taken with the indicated exposure times using an (*A*) 60× or (*B*) 100× objective lens. The color map was scaled to the minimum and maximum pixel values of each image. For the 3-min (*A*) and 5-min (*B*) images, the color bars represent the number of detected photons per pixel. Blue pixels represent saturation of the camera. Fluorescence images (fluo) of cotransfected lifeact-EYFP excited at 491 nm are shown in gray. (Scale bars: 20 μm.)

Overall, no major toxic effects of co *lux* expression were observed. Aldehyde toxicity might not be observed because its concentration is too low. The reason for this could be relatively low aldehyde production by the fatty acid reductase or fast back-conversion of the aldehyde into the acid by the bioluminescence reaction if the activity of all Lux proteins is well-balanced.

### Single-cell bioluminescence imaging

Next, we investigated if the light produced by co *lux* in mammalian cells is sufficient to detect single cells with a custom-built bioluminescence microscope. HEK, HeLa and CV-1 cells were transfected with the co *lux* genes and imaged using different exposure times (Fig. 4). Bioluminescence was already detectable after only 1 s in all cell lines tested and the signal-to-noise ratio increased with prolonged exposure times as expected.

A calibration of the camera demonstrated that several hundred thousand and in some cases more than a million photons per minute per cell were detected.

Since cotransfection of six co *lux* plasmids is required to generate bioluminescence, which might result in a low number of cells expressing all components in sufficient amounts, we determined the fraction of bioluminescent cells. For this purpose, we stained co *lux*-transfected cells with the fluorescent DNA-binding dye Hoechst 33342. This membrane-permeable dye labels the nuclei of living cells and was used to determine the total cell number. Concomitant bioluminescence imaging indicated that ~30% of the cells exhibited detectable bioluminescence emission during the 2-min exposure time and therefore expressed all six genes in sufficient amounts for single-cell imaging (Fig. S6).

We compared the brightness of co *lux* to FLuc and pCMV_Lux_ on the single-cell level. In line with the measurements on whole wells of cells (Figs. 1 and S3), FLuc produced similar levels of bioluminescence as co *lux*, whereas no signal from pCMV_Lux_ was detectable (Fig. S7).

Subsequently, we imaged co *lux*-expressing cells for longer timespans. First, imaging was performed in cell culture medium without additives (Fig. 5*A* and Movie S1). Some cells divided whereas other cells lost their bioluminescence signal, possibly due to cell death resulting from toxicity of the transfection reagent. Since ATP and NADPH are required for sustained bioluminescence, the bioluminescence signal provides information about the metabolic state and cellular viability. As a proof-of-principle application, we tested the influence of the pore-forming toxin gramicidin on co *lux*-expressing cells (Fig. 5*B* and Movie S2). Gramicidin is a peptide antibiotic mainly against Gram-positive bacteria that also affects the ion permeability of eukaryotic cells. It forms ion channels in the cell membrane and thereby disturbs the electrochemical gradient which finally leads to cell death. This became apparent as rounding of the cells and complete loss of bioluminescence within a few hours, confirming its toxicity to eukaryotic cells. The second substance tested was 2,4-dinitrophenol (DNP) that acts as a proton ionophore and thus decouples the proton gradient across the mitochondrial membrane. As a result, ATP synthesis by oxidative phosphorylation is impaired, and the energy of the proton gradient is instead lost as heat. DNP has therefore been used as a drug against obesity, but due to its high toxicity and its severe and sometimes fatal side effects its use is no longer authorized. Up to now, no efficient antidot against DNP poisoning is known. Cells treated with DNP maintained their original shape except for some cases where cells rounded immediately before cell death (Fig. 5*C* and Movie S3), but they moved less on the coverslip than untreated cells (Movie S1). In addition, the signal steadily decreased over time (Fig. 6*A*). A sudden increase in bioluminescence was observed when DNP was removed, but the signal then continued decreasing (Figs. 5*C*, 6*A* and Movie S3). Therefore, the effect of DNP does not seem to be fully reversible, probably either due to its limited water solubility that impedes its washing out or due to causing irreversible cellular damage. Cellular accumulation of DNP might also be responsible for cases of death after repeated intake of sub-lethal DNP doses (28).

**Fig. 5.**
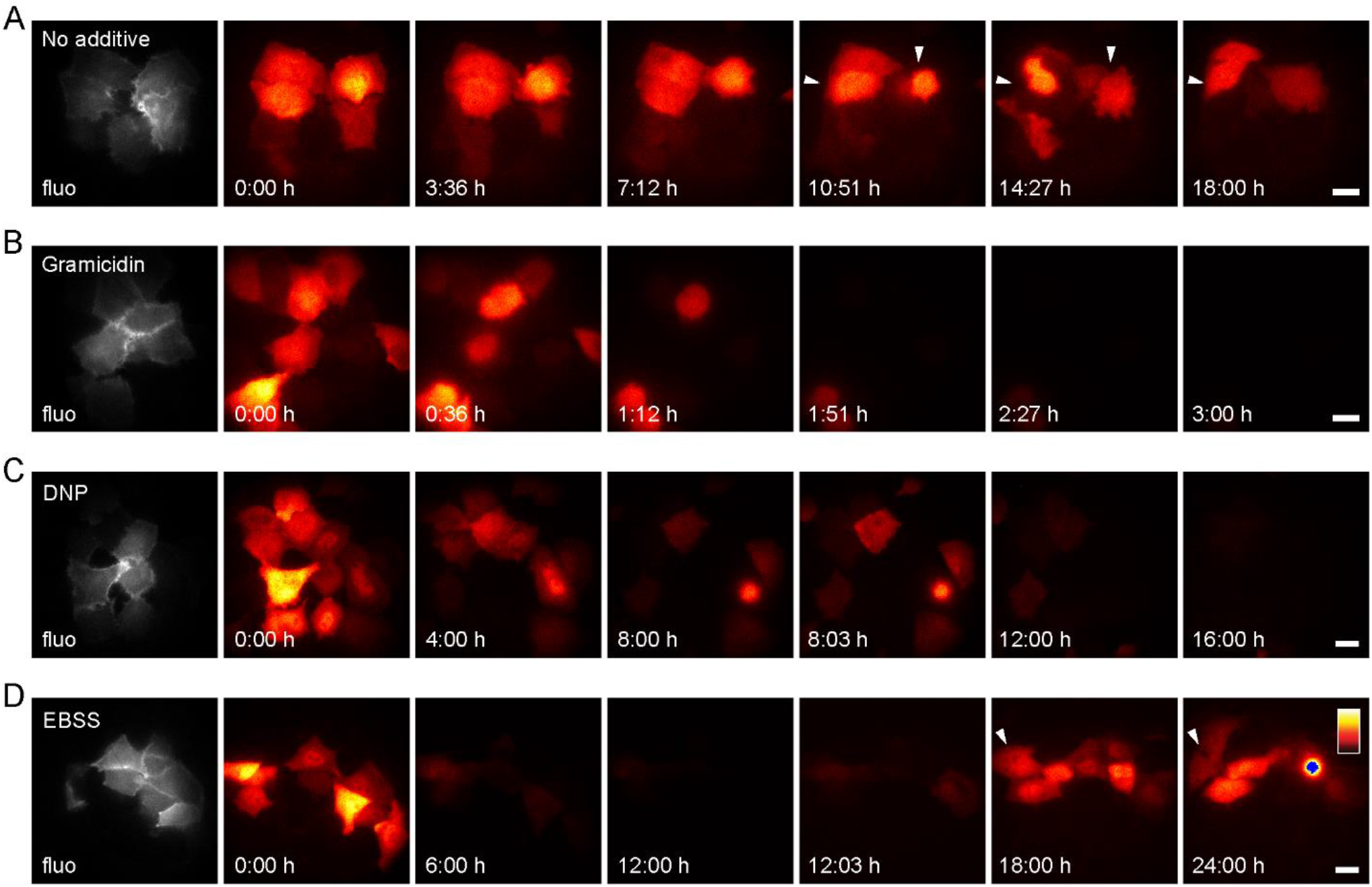
Long-term bioluminescence measurements of HeLa cells expressing co *lux*. Cells were imaged in cell culture medium with (*A*) no additive, (*B*) gramicidin (25 μg/mL) or (*C*) DNP (25 μg/mL) or in EBSS without glucose (*D*). In *C*, cells were washed 5 times with EBSS after 8 h and imaging was continued in cell culture medium without additives. In *D*, cells were washed 10 times with EBSS before starting the measurement, and EBSS was replaced with cell culture medium after 12 h. Bioluminescence images were taken with a 60× objective lens and exposure times of 3 min (*A*, *C*, *D*) or 2 min (*B*). The color map was scaled to the minimum and maximum pixel values of each image series. Blue pixels represent saturation. Arrows indicate dividing cells. Fluorescence images (fluo) of cotransfected lifeact-EYFP excited at 491 nm are shown in gray. Complete time series are shown in Movies S1–S4. (Scale bars: 20 μm.)

Since the energy for the synthesis of ATP and NADPH and thus the generation of bioluminescence is obtained from nutrients in the medium, the bioluminescence signal should decrease upon nutrient deprivation. This effect was investigated by imaging the cells in Earle’s Balanced Salt Solution (EBSS) without glucose instead of cell culture medium (Fig. 5*D* and Movie S4). The signal decreased rapidly during the first ~30 min and then continued decreasing at a lower rate to almost zero after 12 h (Fig. 6*B*). When the EBSS was replaced by cell culture medium, the signal increased immediately and rose further over the next hours (Figs. 5*D*, 6*B* and Movie S4). Although several cells died during the 24-h observation time, other cells survived and divided, demonstrating that some cells are able to recover after the starvation period. The energy source of the prolonged bioluminescence which was still detectable in many cells after 12 h in EBSS is not known. Energy might be derived from remnants of nutrients from the cell culture medium, from intracellularly stored nutrients or uptake of components of the extracellular matrix (29).

**Fig. 6.**
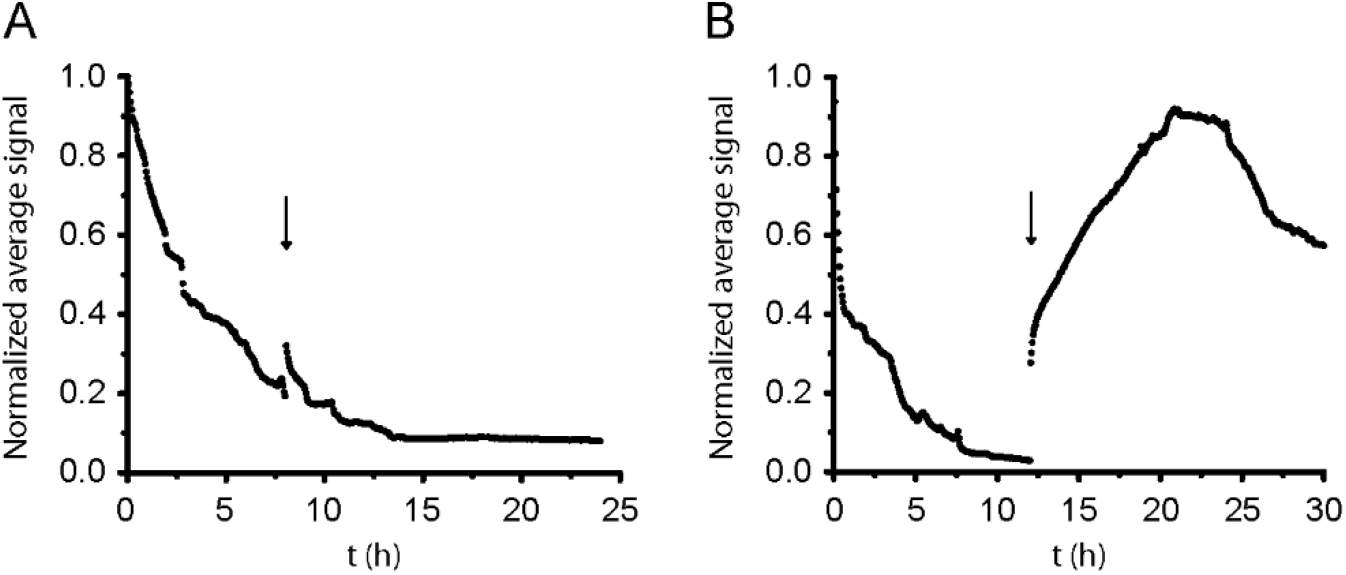
Time course of the bioluminescence signal from HeLa cells expressing co *lux* with DNP and in EBSS. The average signal of the cells in (*A*) Fig. 5*C* (25 μg/mL DNP in cell culture medium) and (*B*) Fig. 5*D* (EBSS without glucose) is shown over the complete time range. Arrows indicate the exchange of the medium or buffer against fresh cell culture medium without additives.

As we previously reported for *E. coli* cells (30), the bioluminescence signal from a few HeLa cells fluctuated before cell death (Movies S3 and 4). This “blinking” occurred less frequently than in kanamycin-treated *E. coli* cells where it was shown to be accompanied by changes in the cellular ATP concentration (30). The cause of the blinking is unknown, but might involve similar processes affecting the energy metabolism during cell death in both cell types.

### Discussion and Conclusion

We demonstrated for the first time autonomous bioluminescence imaging of single mammalian cells by coexpression of six codon-optimized genes from the bacterial bioluminescence system. Although the brightness of bacterial bioluminescence has so far been considered to be much lower than that of other luciferases requiring an external luciferin, we found that co Lux produced autonomous bioluminescence at levels similar to firefly luciferase. Codon optimization was found to be crucial for high expression in mammalian cells, although we cannot rule out that features of the nucleotide sequence other than the codon usage contribute to enhanced expression (e.g., due to altered binding of transcription factors or formation of mRNA secondary structures). In addition, several other factors affect the expression or function of the Lux proteins and may explain the previously observed decreased performance of bacterial bioluminescence in mammalian cells: (i) choice of the specific LuxCDABE and flavin reductase amino acid sequences, as the proteins from different bacterial species may differ in their stability at 37 °C and their expression, maturation or activity under the conditions in mammalian cells, which we observed in particular for the flavin reductase, (ii) the presence of fusion partners at the N and C terminus of the Lux proteins, as even the small 2A peptides can strongly reduce the bioluminescence signal, (iii) the ratio at which the six genes are transfected, since aldehyde synthesis seems to be the most rate-limiting step of the overall process and expression of all co *lux* genes from individual plasmids allows to freely adjust the protein ratios, and (iv) choice of the expression vector and distance of the *lux* genes from the promoter.

Although the aldehyde produced by the Lux system may be cytotoxic for eukaryotic cells (31), we did not observe significant effects on cell number or metabolic activity 24 h post-transfection, which is in line with other reports (11, 12, 32). Furthermore, co *lux*-expressing cells maintained their physiological shape and their ability to divide. The only significant effect of co *lux* expression was found to be a small decrease of NADPH levels which possibly results from elevated NADPH consumption during synthesis of the aldehyde or FMNH_2_. It is not clear whether this represents already the metabolic limit for the light levels achievable by autonomous bioluminescence due to its high energy demand, or if the brightness could be further increased by optimization of the involved proteins and their expression levels in mammalian cells.

Because of its dependence on metabolic energy, bioluminescence generated by the co lux genes can be used to investigate metabolic changes in mammalian cells, e.g. during the cell cycle phases, during nutrient deprivation or in response to toxic compounds. On the other hand, the generation of bioluminescence requires substantial amounts of metabolic energy, which may itself influence cellular processes. Bioluminescence measurements with co Lux can easily be performed with cell monolayers using a commercial luminescence imager. Owing to the high bioluminescence signal, they are also feasible on the single-cell level which allows the observation of individual differences between cells and events such as fluctuations in brightness, cell division and cell death. Therefore, co Lux holds promise as an easy-to-use reporter system for numerous applications in biological and medical research. Although fluorescence-based methods remain superior for imaging of cells at high speed and resolution due to their much higher signals, further improvements of the co Lux system will extend its scope of application in singlecell imaging.

## Materials and Methods

Details of plasmid generation, cell culture, bioluminescence imaging and toxicity tests are described in SI Materials and Methods. In brief, co *lux* and *frp* genes were cloned separately into the vector pcDNA3.1(+) and cotransfected into mammalian cell lines. Bioluminescence imaging was performed with a custom microscope containing a cooled electron multiplying charge-coupled device (EMCCD) camera and a 491-nm excitation laser for fluorescence measurements.

## Acknowledgements

We thank Rainer Pick and Dr. Ellen Rothermel for technical assistance. We thank Dr. Steven Ripp (490 BioTech) for kindly providing the pCMV_Lux_ plasmid and Prof. Yi Yang (East China University of Science and Technology) for the iNap1 pcDNA3.1/Hygro(+) plasmid.

## Author Contributions

C.G. and S.W.H. designed research; C.G. performed research; J.P and K.C.G. built the microscope; T.G. performed cell culture and transfections; C.G. analyzed data; C.G., S.J.S. and S.W.H. wrote the paper.

## Supplementary Information

### SI Materials and Methods

#### Codon optimization of the *lux* genes and cloning

The *luxC*, *D*, *A*, *B* and *E* genes from *Photorhabdus luminescens* (GenBank accession number JXSK01000001.1) and the *frp* gene from *Vibrio campbellii* (GenBank accession number AGU98260.1) were codon-optimized (co) for expression in human cells with the codon adaptation tool JCat (http://www.jcat.de, (1)). The resulting genes (Fig. S1) were synthesized (Synbio Technologies) and cloned into the expression vector pcDNA3.1(+) between the NheI and XhoI restriction site (BamHI and XhoI for co *lux*E). Wild-type *lux* genes without codon optimization were cloned into pcDNA3.1(+) between the BamHI and XhoI restriction site. Firefly luciferase (FLuc, GenBank accession number MK484107.1) and h*lux* genes were cloned into pcDNA3.1(+) between the NheI and XhoI restriction site.

co *luxAB*, co *luxBA*, co *luxCD* and co *luxDE* were generated by integration of a GSG linker and a P2A sequence between the two genes (GSGATNFSLLKQAGDVEENPGP, nucleotide sequence GGATCCGGCGCCACCAACTTCAGCCTGCTGAAGCAGGCCGGCGACGTGGAGGAGAACCCCGGCCCC). For construction of co *luxABfrp* and co *luxDEC*, the same P2A sequence was inserted between the first and second gene and a T2A sequence (GSGEGRGSLLTCGDVEENPGP, nucleotide sequence GGCAGCGGCGAGGGCCGCGGCAGCCTGCTGACCTGCGGCGACGTGGAGGAGAACCCCGGGCCC) between the second and third gene. The complete inserts were cloned into pcDNA3.1(+) between the NheI and XhoI restriction site.

pRsetB-PercevalHR was a gift from Gary Yellen (Addgene plasmid #49081). pC1-HyPer-3 was a gift from Vsevolod Belousov (Addgene plasmid #42131). PercevalHR and HyPer-3 were amplified by PCR and cloned into pcDNA3.1(+) between the NheI and XhoI restriction site.

##### Cell culture

All cell lines were cultured in DMEM containing 4.5 g/l glucose without phenol red (Gibco) supplemented with 10% FBS (Biochrom), 1 mM sodium pyruvate (Merck), 100 units/mL penicillin and 100 μg/mL streptomycin (Biochrom). Cells were cultured at 37 °C in 5% CO_2_.

##### Bioluminescence imaging

Comparison of the bioluminescence signal from different constructs was performed with an Amersham Imager 600 (GE Healthcare). Cells were seeded in 24-well plates with 500 μL of cell culture medium per well without coverslips and transfected with a total amount of 500 ng DNA using 1 μL of Lipofectamine 2000 (Invitrogen) per well according to the protocol of the manufacturer. Imaging was performed 24 h post-transfection. For FLuc, D-luciferin (Merck) was added to the medium at a concentration of 150 μg/mL 5 min prior to imaging. Coelenterazine (Carl Roth) was used at a concentration of 5 μM.

For single-cell imaging, cells were seeded in 12-well plates on 18 mm round coverslips with 1 mL of cell culture medium per well. Cells were transfected with a mixture of the following plasmids using 2 μL of Lipofectamine 2000: 83 ng co *luxA*, 83 ng co *luxB*, 250 ng co *luxC*, 250 ng co *luxD*, 250 ng co *luxE*, 83 ng co *frp* and 50 ng lifeact-EYFP (all in pcDNA3.1(+)). Cells were imaged the following day with a custom-built setup described in (2) with the following modifications: The electron multiplying charge-coupled device (EMCCD) camera was exchanged with an iXon EMCCD DU-897E-CS0-#BV (Andor) for a larger field of view (512×512 pixels). The camera sensor was cooled to –100 °C. An oil immersion objective lens (HC PL APO CS 100×/1.40–0.70 OIL (Leica) or PlanApo N 60×/1.42 (Olympus)) was used to collect the emitted light. The lens focusing on the camera was exchanged to reduce the focal length (AC254-125-A-ML; Thorlabs), resulting in an increased effective pixel size on the camera of 230 nm for the 100× and 360 nm for the 60× objective. The dielectric mirrors were replaced by mirrors with a diameter of 2” (BB2-E02; Thorlabs). Cells were focused by their lifeact-EYFP fluorescence excited with a 491-nm laser (Calypso 50 mW; Cobolt). Imaging was performed in a humidified chamber in 5% CO_2_ at 37 °C (CO2-UNIT-BL and H301-T-UNIT-BL-PLUS; Okolab) without the focus lock system. Calibration of camera pixel counts to detected photons was performed as described in (2). Bright pixels resulting from cosmic rays were filtered out using a custom-written Matlab (MathWorks) script by comparison of each pixel value either to the surrounding pixels or the same pixel in the previous and following image and replacement by their average if the difference was above a threshold value. The color map was then scaled to the minimum and maximum pixel values of each image or image series, respectively.

To determine the fraction of cells that were transfected with all six co *lux* plasmids, HeLa cells grown on 18 mm coverslips were transfected with a mixture of the co *lux* pcDNA3.1(+) plasmids as described above. 24 h post-transfection, cells were stained with Hoechst 33342 (Molecular Probes) at a concentration of 10 μg/mL in cell culture medium for 5 min. Subsequently, cells were washed and imaged in cell culture medium as described above. Hoechst fluorescence was excited at 405 nm, bioluminescence images were taken with an exposure time of 1–2 min. The total cell number was determined by counting of the Hoechst-labeled nuclei in 10 regions (corresponding to ~150 cells in total) of each coverslip. The number of bioluminescent cells was determined from the bioluminescence images of the same regions that were subsequently taken. The fraction of bioluminescent cells was accordingly calculated as the number of bioluminescent cells divided by the total cell number for each coverslip.

Gramicidin and 2,4-dinitrophenol (DNP) were purchased from Sigma-Aldrich. EBSS (Earle’s Balanced Salt Solution) without glucose contained 200 mg/L CaCl_2_, 200 mg/L MgSO_4_·7 H_2_O, 400 mg/L KCl, 2,2 g/L NaHCO_3_, 6,8 g/L NaCl and 140 mg/L NaH_2_PO_4_. The pH was adjusted to 7.0 at 37 °C and 5% CO_2_.

##### Toxicity control tests

All cells were grown on 18 mm coverslips in 1 mL of cell culture medium and transfected with 2 μL of Lipofectamine 2000.

For the MTT assay, cells were transfected with 1 μg of empty pcDNA3.1(+) vector or a mixture of 83 ng co *luxA*, 83 ng co *luxB*, 250 ng co *luxC*, 250 ng co *luxD*, 250 ng co *luxE* and 83 ng co *frp* (all in pcDNA3.1(+)). 24 h post-transfection, 100 μL of MTT (Sigma-Aldrich) in PBS (5 mg/mL) was added. After incubation at 37 °C in 5% CO_2_ for 1 h, 1 mL 10% SDS was added and cells were resuspended by pipetting. Absorbance at 570 nm was measured with a NanoDrop 1000 Spectrophotometer (Peqlab Biotechnologie GmbH).

Intracellular ATP, NADPH and H_2_O_2_ concentrations were compared by fluorescence imaging with the PercevalHR, iNap1 and HyPer-3 sensor, respectively. Cells were cotransfected with 200 ng PercevalHR pcDNA3.1(+), iNap1 pcDNA3.1/Hygro(+) or HyPer-3 pcDNA3.1(+), respectively, and 800 ng empty pcDNA3.1(+) vector or a mixture of 67 ng co *luxA*, 67 ng co *luxB*, 200 ng co *luxC*, 200 ng co *luxD*, 200 ng co *luxE* and 67 ng co *frp* (all in pcDNA3.1(+)). 24 h post-transfection, cells were imaged at a confocal SP8 microscope (Leica). Imaging was performed with a pixel size of 480 nm and a detection range of 510–570 nm. Identical laser powers were used for excitation at the indicated wavelength for all images of the respective sensor. The fluorescence ratio for a given cell was calculated by dividing the total signals in both channels. p-values of the fluorescence ratios of 50 cells transfected with co *lux* or empty pcDNA3.1(+) vector were calculated by a two-tailed Student’s *t*-test.

**Fig. S1.**
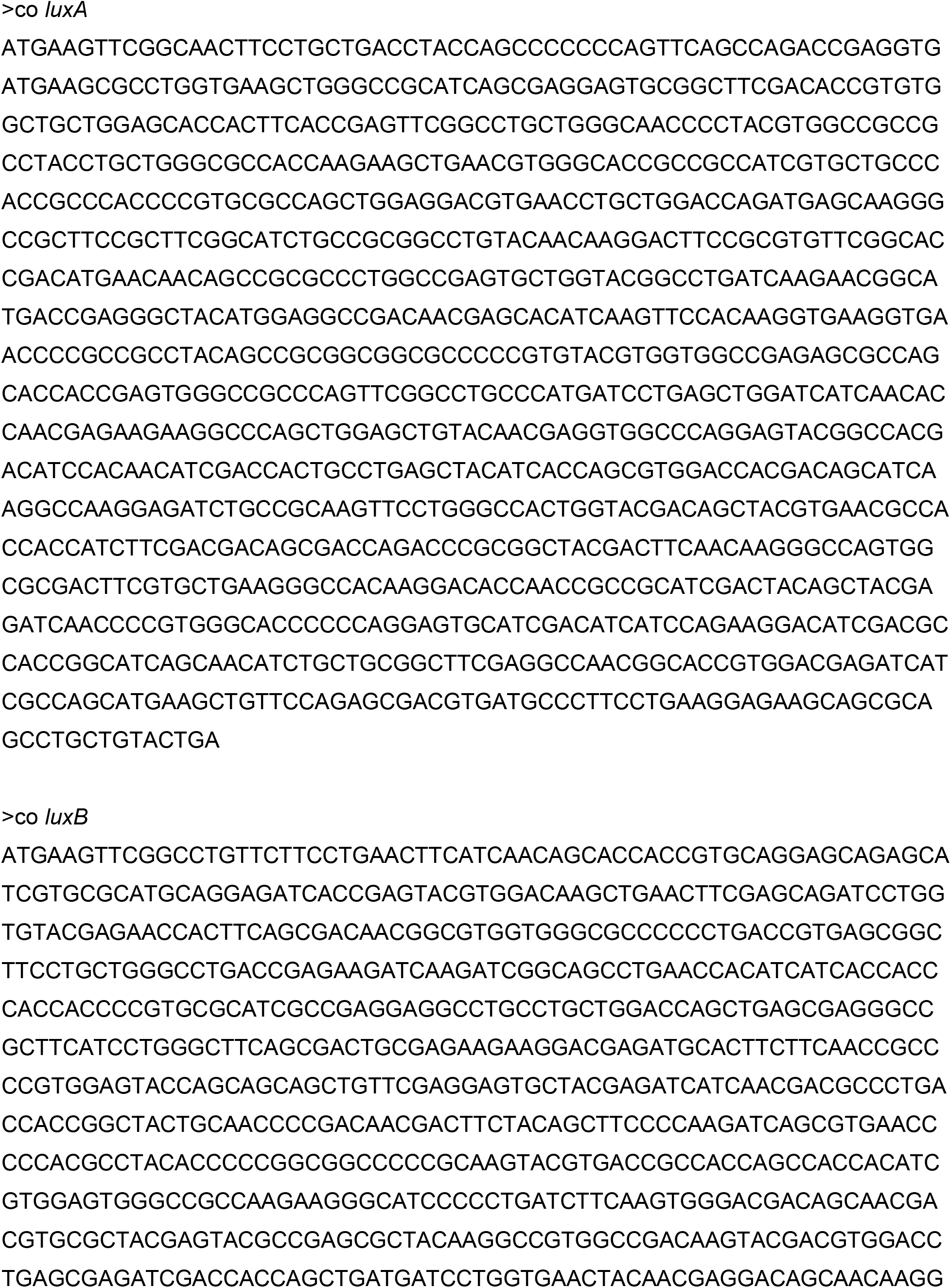
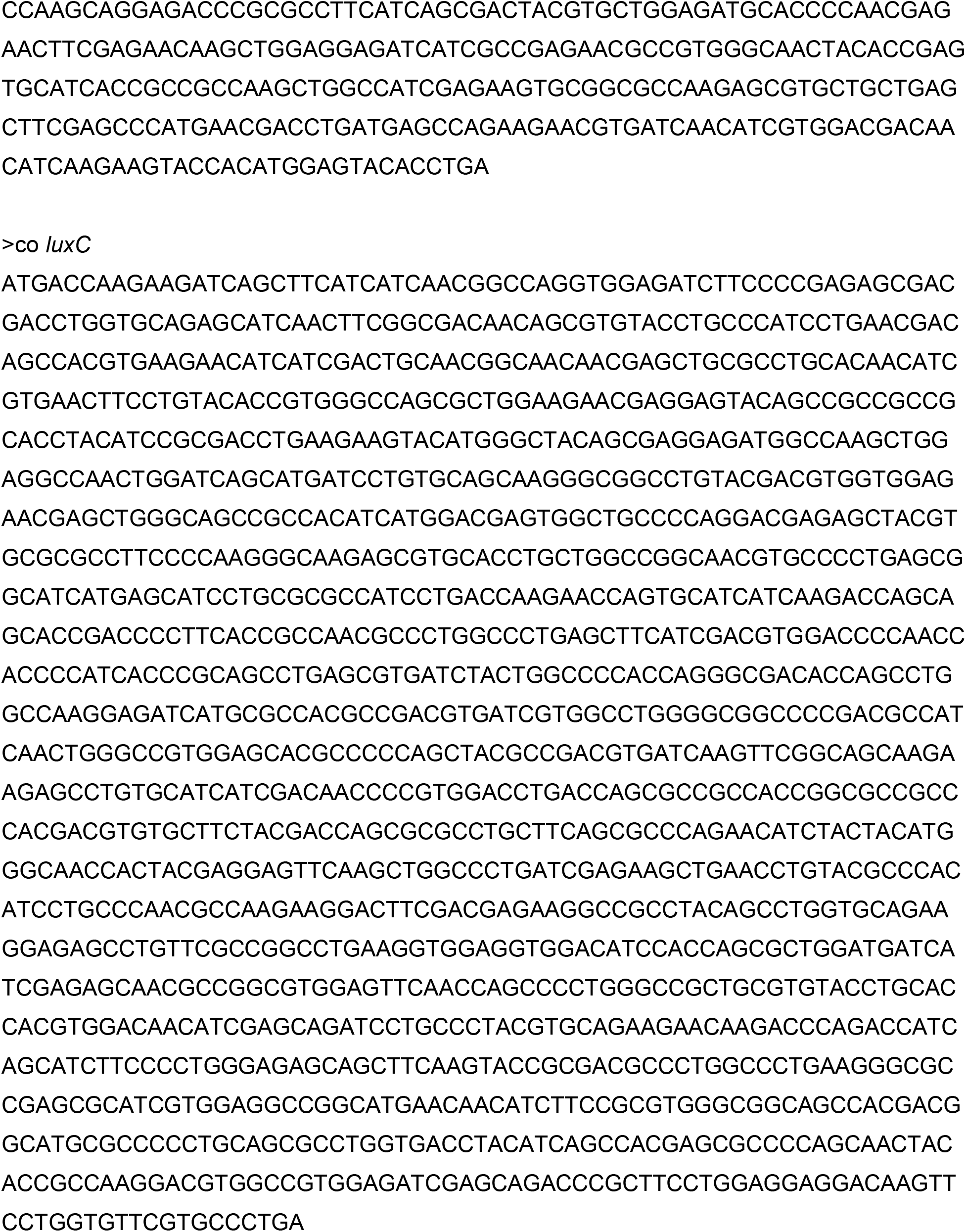
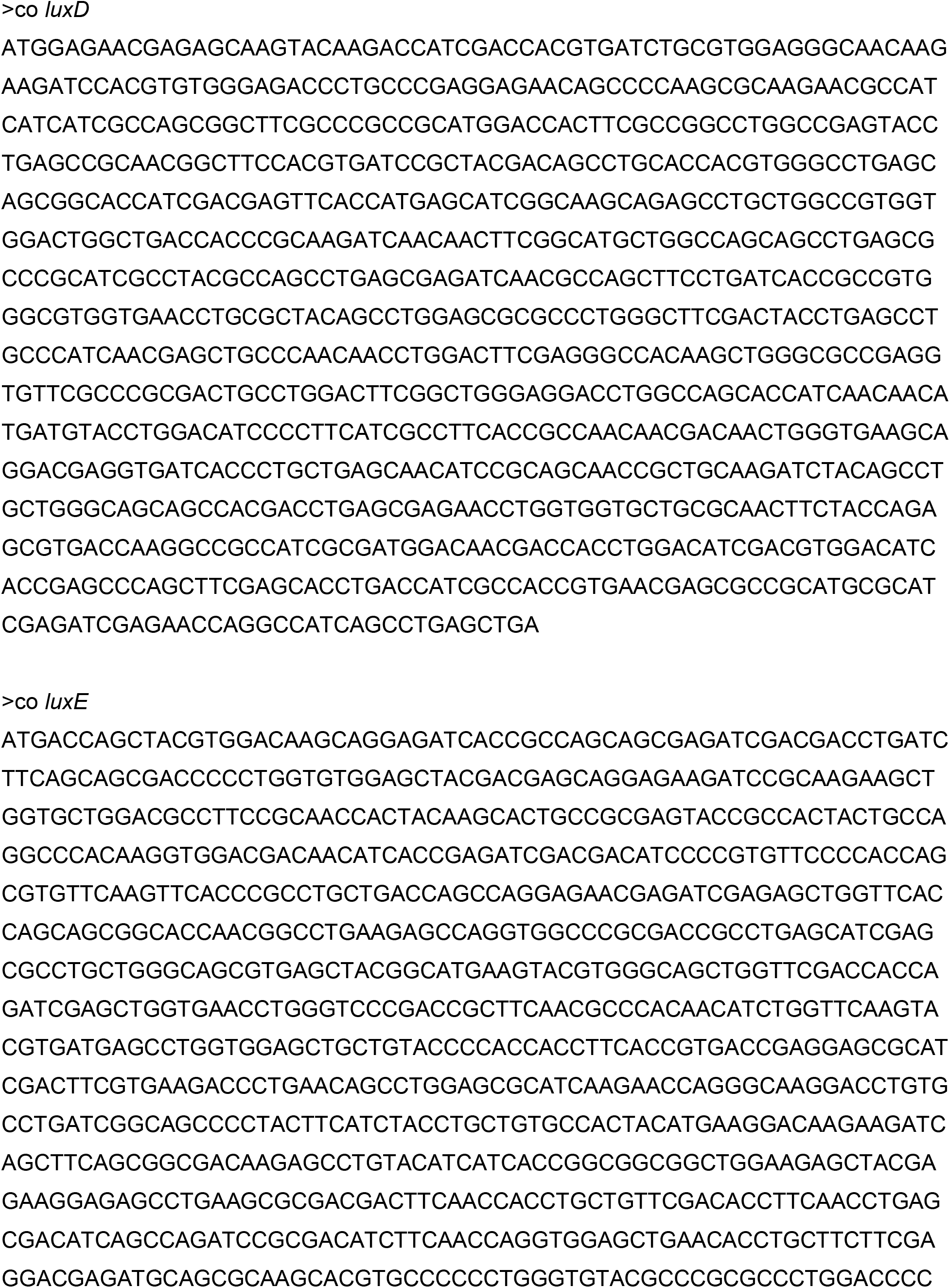
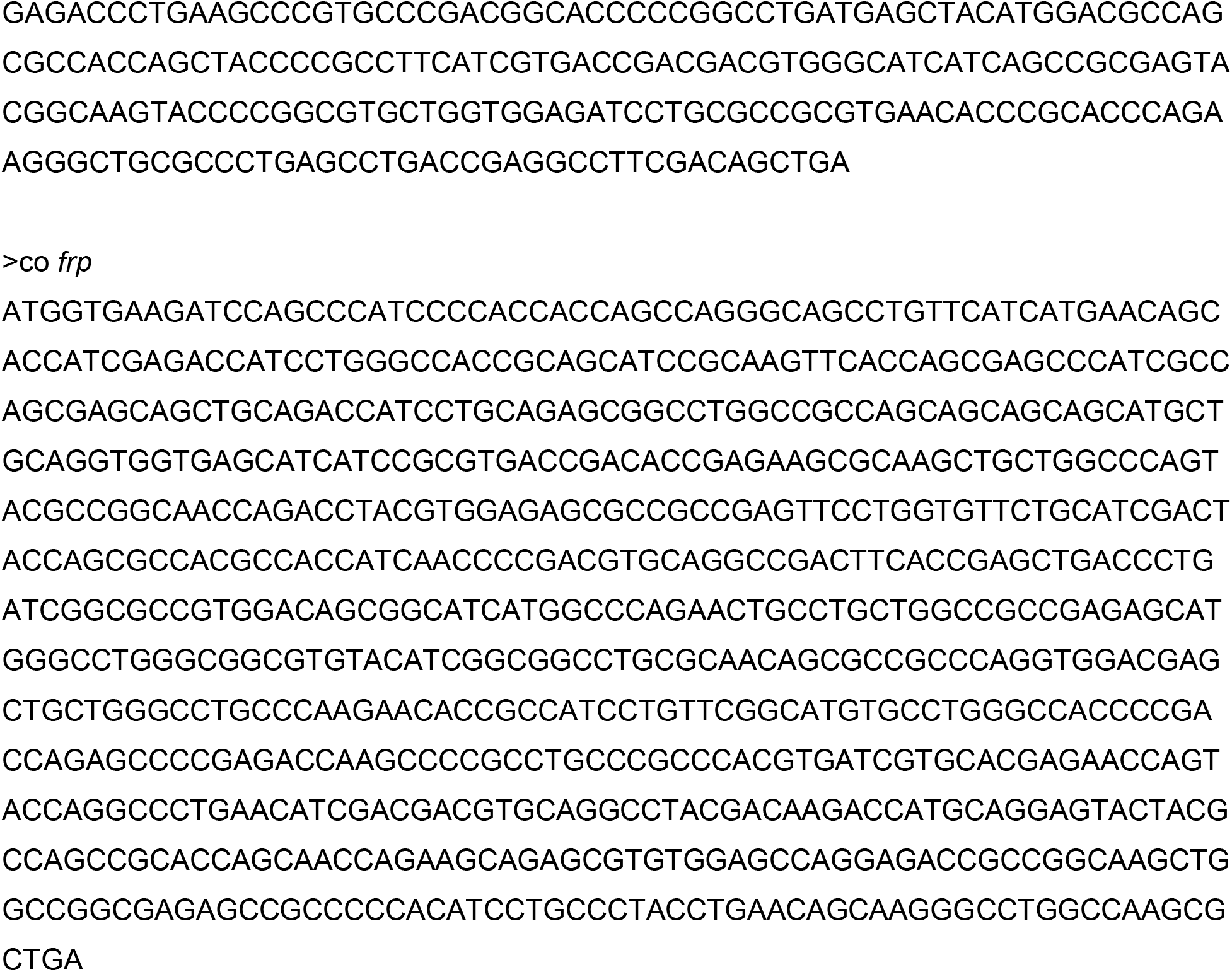
Nucleotide sequences of the codon-optimized (co) *lux* genes.

**Fig. S2.**
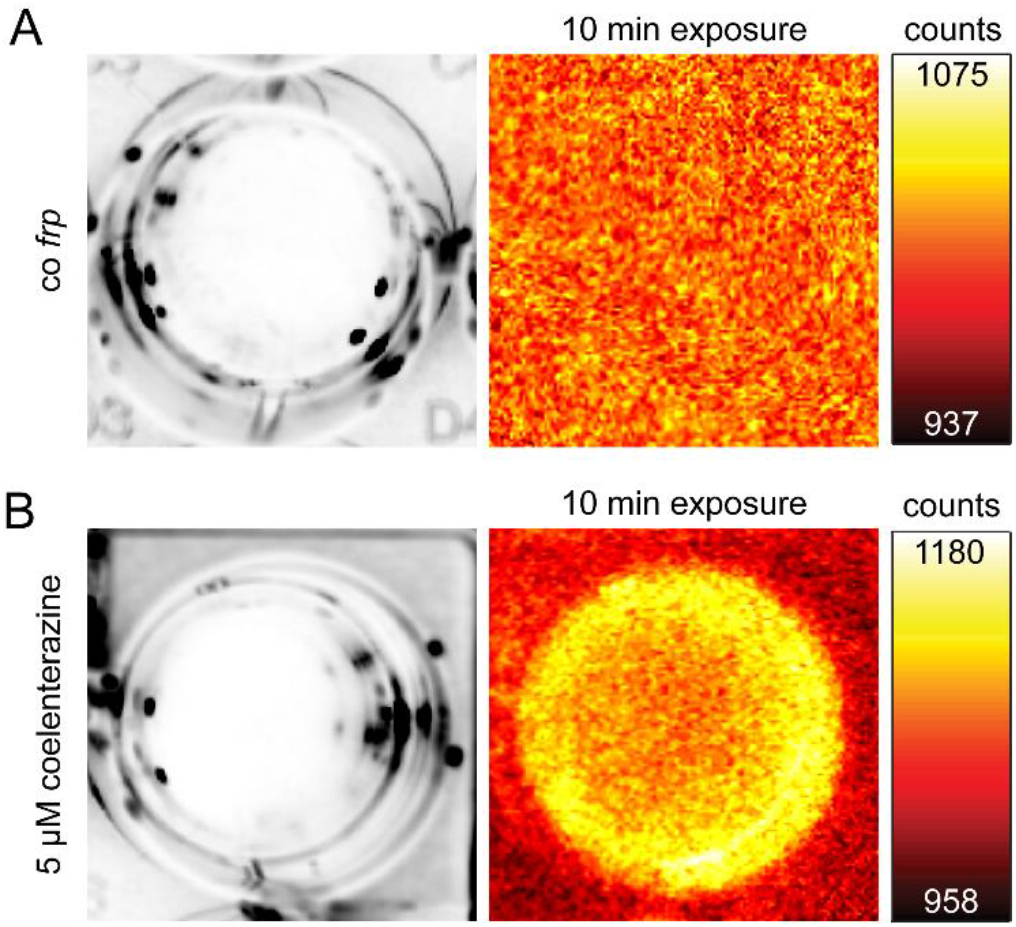
Comparison of background signal from coelenterazine and FMNH_2_. HEK 293 cells were transfected with 42 ng co *frp* pcDNA3.1(+) and 458 ng empty pcDNA3.1(+) vector (*A*) or not transfected and 5 μM coelenterazine added before imaging (*B*). Cells were imaged in a 24-well plate with an Amersham Imager 600. Left, white-light images of the respective well with a monolayer of cells, right, bioluminescence signal after an exposure time of 10 min. The color maps were scaled to the minimum and maximum camera counts per pixel of the bioluminescence images.

**Fig. S3.**
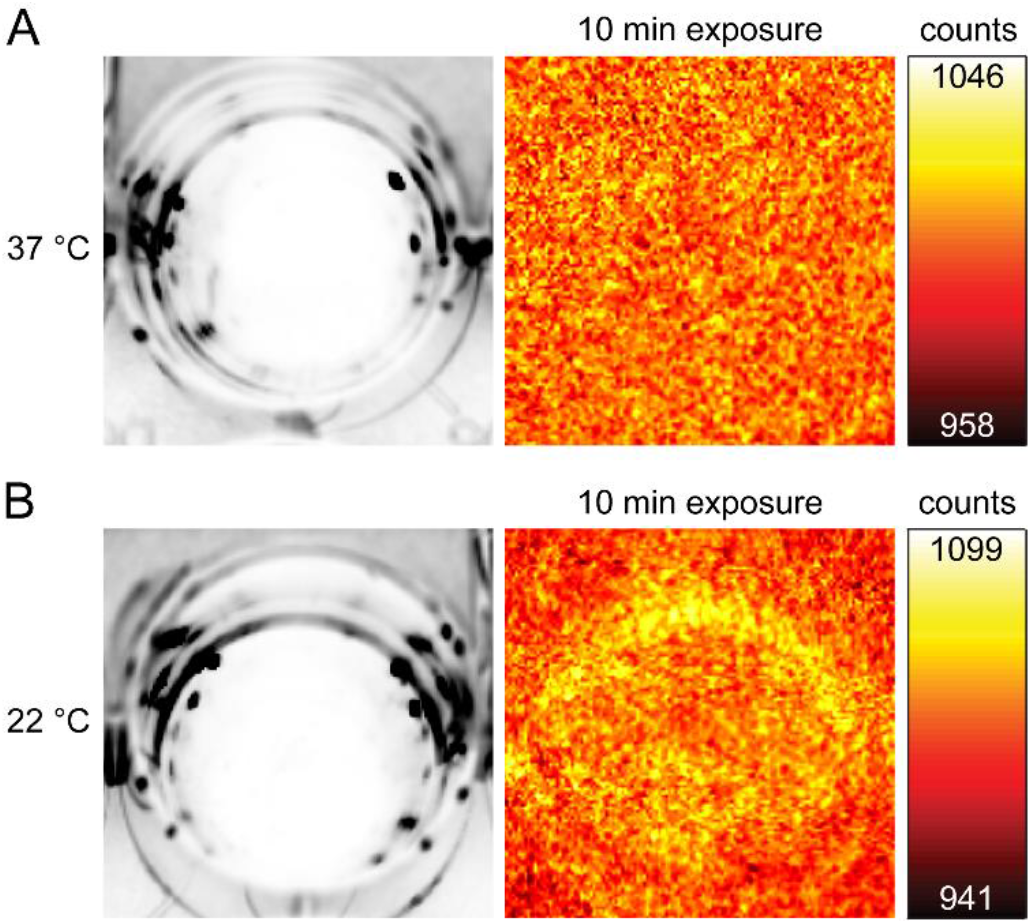
Bioluminescence signal from pCMV_Lux_. HEK 293 cells were transfected with 500 ng pCMV_Lux_ and imaged at 37 °C (*A*) or 22 °C (*B*). Cells were imaged in a 24-well plate with an Amersham Imager 600. Left, white-light images of the respective well with a monolayer of cells, right, bioluminescence signal after an exposure time of 10 min. The color maps were scaled to the minimum and maximum camera counts per pixel of the bioluminescence images.

**Fig. S4.**
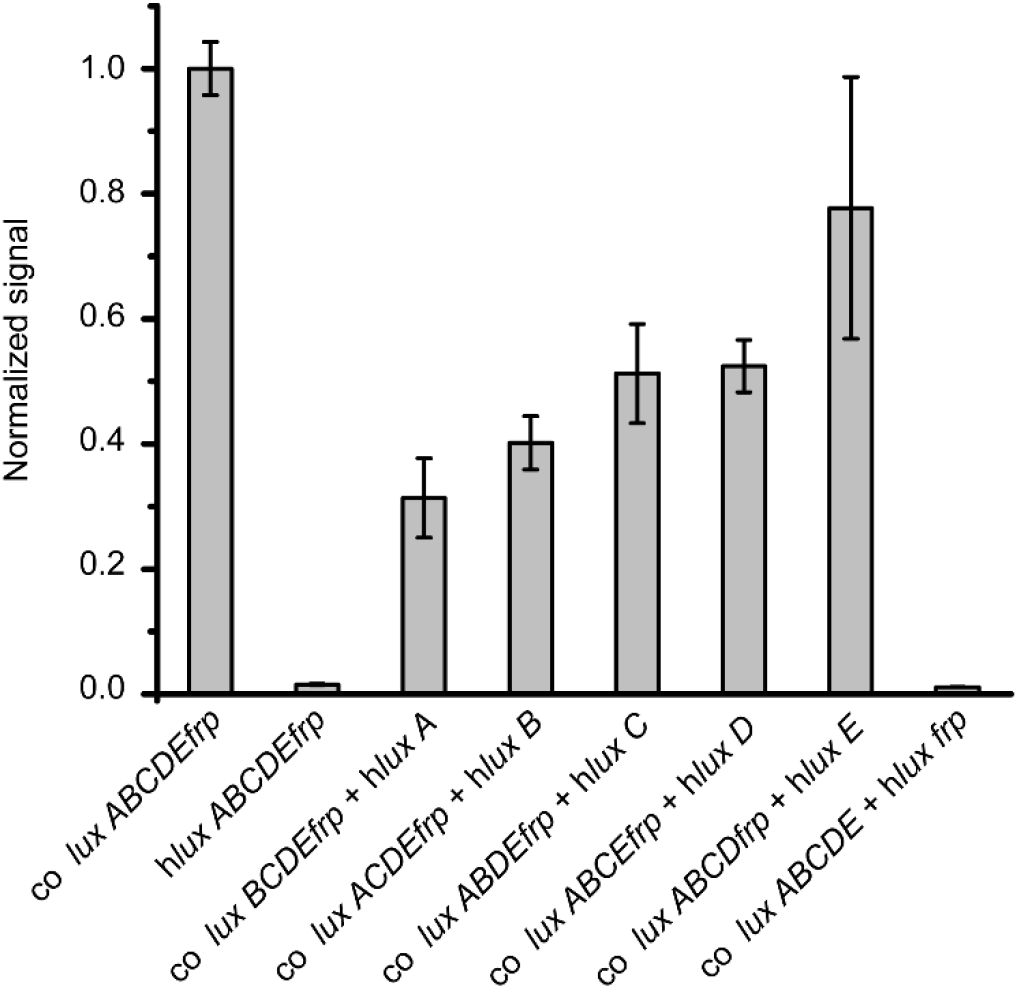
Bioluminescence of HEK 293 cells expressing different combinations of co *lux* and h*lux* genes from pcDNA3.1(+). Cells were grown in 24-well plates, transfected with the same total amount of DNA and imaged with an Amersham Imager 600. Plasmids encoding *luxA*, *B*, *C*, *D*, *E* and *frp* were transfected in a ratio of 1:1:3:3:3:1. Error bars represent SD of the signal of five separate wells.

**Fig. S5.**
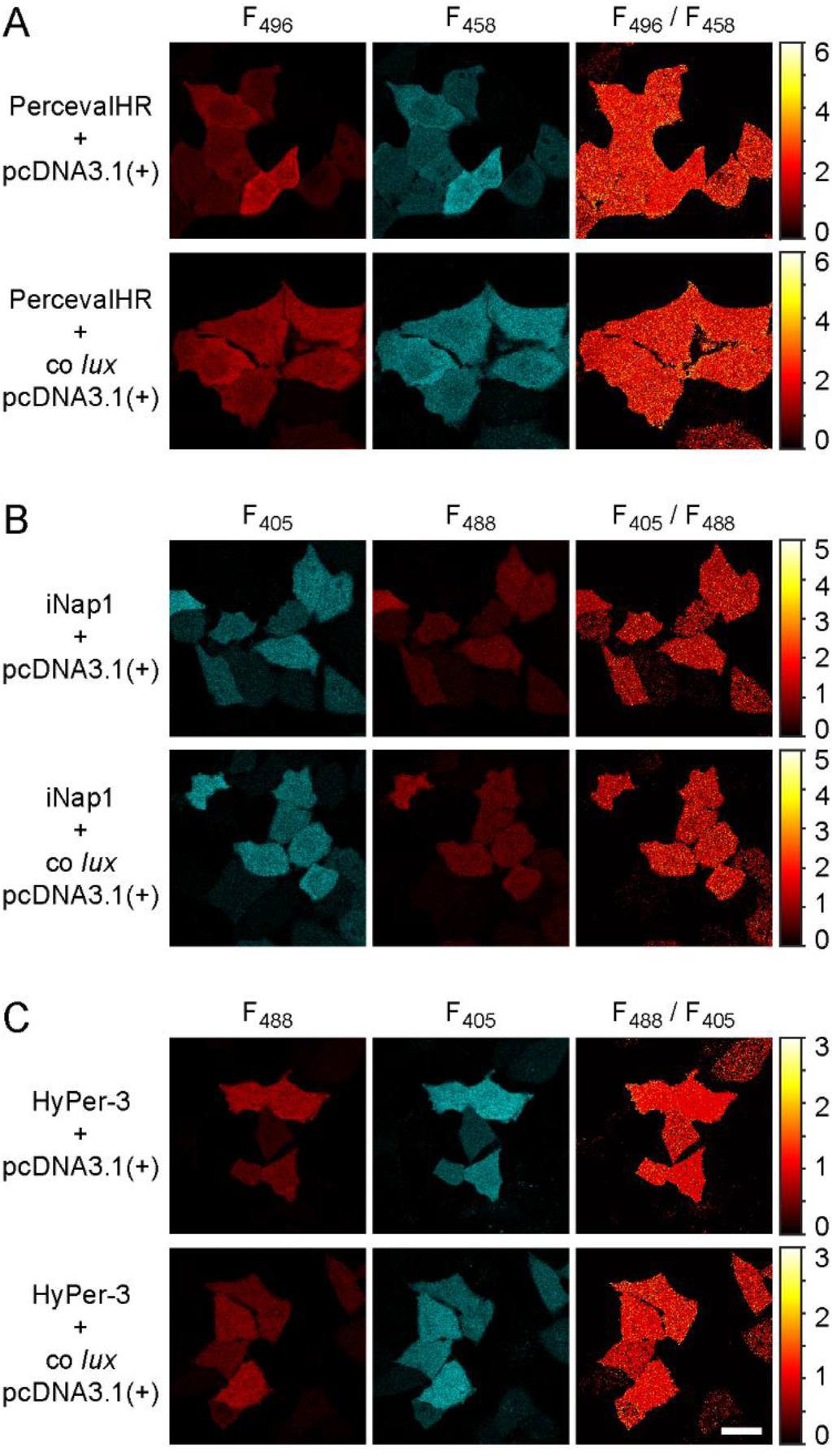
Comparison of intracellular ATP, NADPH and H_2_O_2_ levels in HeLa cells cotransfected with the respective fluorescent sensor and empty pcDNA3.1(+) vector or co *lux* genes in pcDNA3.1(+). Representative images of the ATP sensor PercevalHR (*A*), the NADPH sensor iNap1 (*B*) and the H2O2 sensor HyPer-3 (*C*) are shown. The colorbar represents the fluorescence ratio at the indicated excitation wavelengths. Blue pixels represent saturation of the color map. (Scale bar: 50 μm.)

**Fig. S6.**
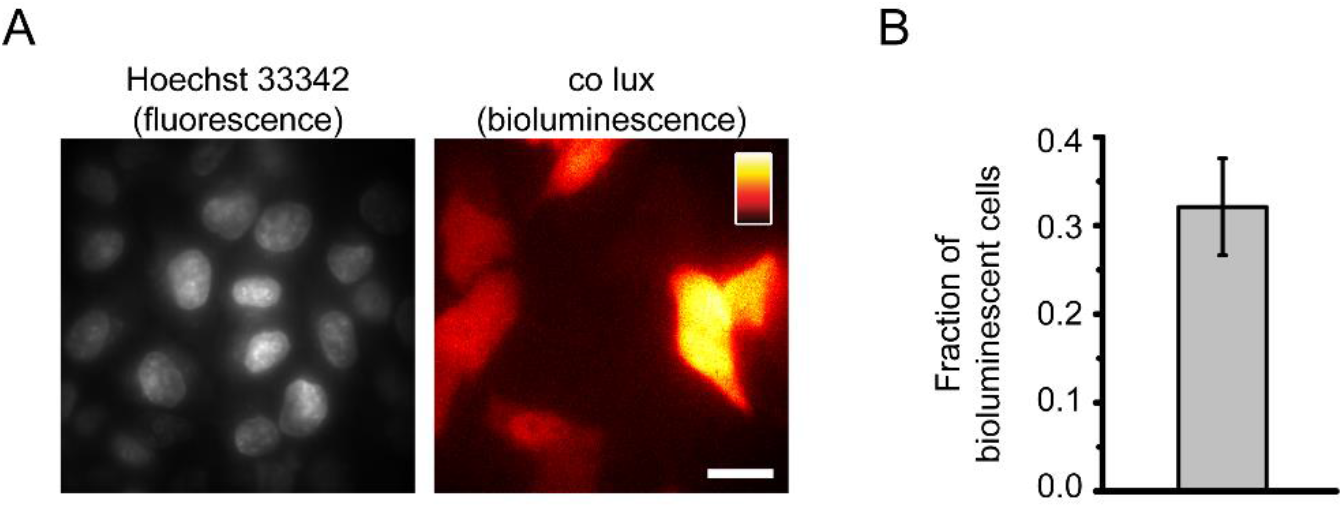
Determination of the fraction of co *lux*-expressing cells. HeLa cells transfected with co *lux* were stained with Hoechst 33342. (*A*) Images of Hoechst fluorescence excited at 405 nm (left) and bioluminescence taken with a 2-min exposure time (right) using a 60× objective lens. The color maps were scaled to the minimum and maximum pixel values of each image. (Scale bar: 20 μm.) (*B*) Fraction of bioluminescent cells. Error bars represent SD from four separate coverslips.

**Fig. S7.**
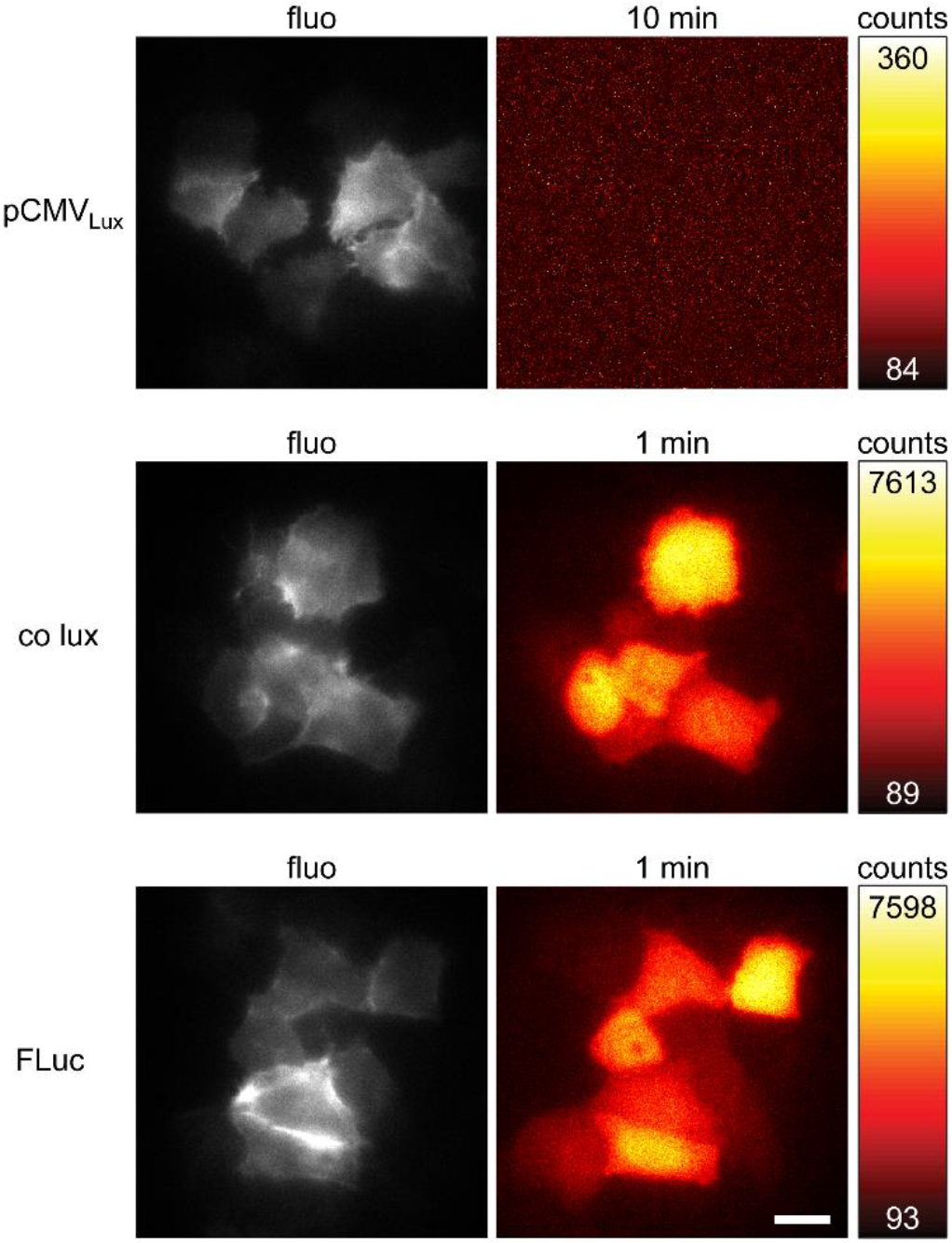
Comparison of bioluminescence from pCMV_Lux_, co lux and FLuc. HeLa cells were transfected with 1 μg of pCMV_Lux_, co *lux* mixture or FLuc. For FLuc, 150 μg/mL D-luciferin was added to the medium prior to imaging. Fluorescence images (fluo, left) of cotransfected lifeact-EYFP excited at 491 nm are shown in gray. Bioluminescence images (right) were taken with the indicated exposure times. The color maps of the bioluminescence images were scaled to the minimum and maximum pixel values which are indicated in the color bar. (Scale bar: 20 μm.)

**Table S1.**
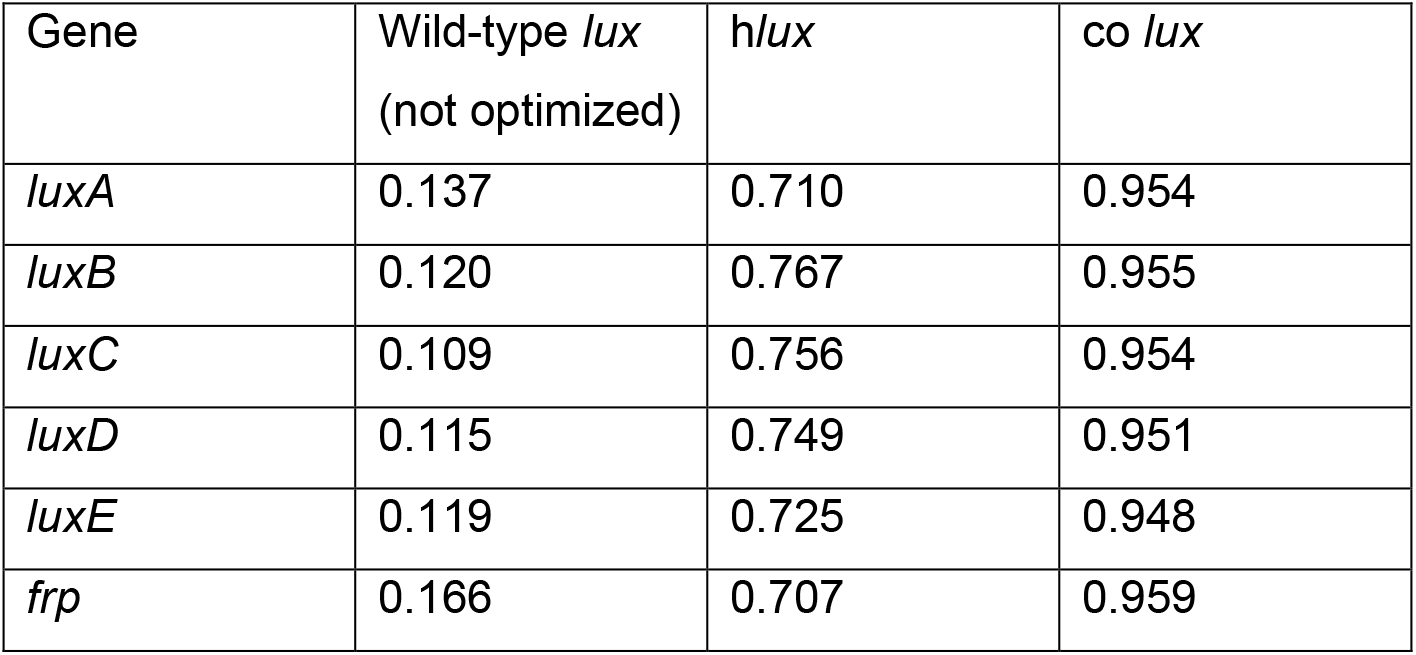
Comparison of codon adaptation index (CAI) of the *lux* genes. Values were calculated with JCat (http://www.jcat.de, (1)) for *Homo sapiens*.

### Supplementary Movie Legends

**Movie S1.** Cell division of HeLa cells expressing co *lux*. Cells were imaged in cell culture medium without additives. Single images were taken with 3-min exposure time. (Scale bar: 20 μm.)

**Movie S2.** HeLa cells expressing co *lux* treated with gramicidin. Cells were imaged in cell culture medium with gramicidin (25 μg/mL). Single images were taken with 2-min exposure time. (Scale bar: 20 μm.)

**Movie S3.** HeLa cells expressing co *lux* treated with DNP. Cells were imaged in cell culture medium with DNP (25 μg/mL). After 8 h, cells were washed and the medium was replaced by fresh cell culture medium without additives. Single images were taken with 3-min exposure time. (Scale bar: 20 μm.)

**Movie S4.** HeLa cells expressing co *lux* imaged in EBSS without glucose. After 12 h, EBSS was replaced by fresh cell culture medium without additives. Single images were taken with 3-min exposure time. (Scale bar: 20 μm.)

